# Efficient identification of trait-associated loss-of-function variants in the UK Biobank cohort by exome-sequencing based genotype imputation

**DOI:** 10.1101/2021.08.12.456052

**Authors:** Lei Zhang, Shan-Shan Yan, Jing-Jing Ni, Yu-Fang Pei

## Abstract

The large-scale open access whole-exome sequencing (WES) data of the UK Biobank ~200,000 participants is accelerating a new wave of genetic association studies aiming to identify rare and functional loss-of-function (LoF) variants associated with a broad range of complex traits and diseases, however the community is in short of stringent replication of new associations. In this study, we proposed to merge the WES genotypes and the genome-wide genotyping (GWAS) genotypes of 167,000 UKB Caucasian participants into a combined reference panel, and then to impute 241,911 UKB Caucasian participants who had the GWAS genotypes only. We then proposed to use the imputed data to replicate association identified in the discovery WES sample. Using a leave-100-out imputation strategy in the reference panel, we showed that average imputation accuracy measure r^2^ is modest to high at LoF variants of all minor allele frequency (MAF) intervals including ultra-rare ones: 0.942 at MAF interval [1%, 50%], 0.807 at [0.1%, 1.0%), 0.805 at [0.01%, 0.1%), 0.664 at [0.001%, 0.01%) and 0.410 at (0, 0.001%). As applications, we studied single variant level and gene level associations of LoF variants with estimated heel BMD (eBMD) and 4 lipid traits: high-density-lipoprotein cholesterol (HDL-C), low-density-lipoprotein cholesterol (LDL-C), triglycerides (TG) and total cholesterol (TC). In addition to replicating dozens of previously reported genes such as *MEPE* for eBMD and *PCSK9* for more than one lipid trait, the results also identified 2 novel gene-level associations: *PLIN1* (cumulative MAF=0.10%, discovery BETA=0.38, P=1.20×10^−13^; replication BETA=0.25, P=1.03×10^−6^) and *ANGPTL3* (cumulative MAF=0.10%, discovery BETA=−0.36, P=4.70×10^−11^; replication BETA=−0.30, P=6.60×10^−11^) for HDL-C, as well as one novel single variant level association (11:14843853:C:T, MAF=0.11%, discovery BETA=−0.31, P=2.70×10^−9^; replication BETA=−0.31, P=8.80×10^−14^, *PDE3B*) for TG. Our results highlighted the strength of WES based genotype imputation as well as provided useful imputed data within the UKB cohort.

## Introduction

Whole-exome sequencing (WES) technology is an efficient approach in capturing coding genomic variation. The development of next-generation sequencing technologies and sequence/target enrichment methods has made WES both technically feasible and cost-effective, emerging large-scale WES studies at a hundreds of thousands participants scale [1, 2]. These studies, equipped with sufficient sample size, have unraveled potential pathological significance of rare coding variation to human health. Among various types of coding variants, loss-of-function (LoF) variants, due to their severe and interpretable relevance to personal health, are the center of research focus and clinical application.

With the release of the WES data for an interim of ~50,000 participants and then ~150,000 participants, the UKB is now providing a largest open access WES genomic resource of coding variants in ~200,000 participants [3]. With no doubt, these data will accelerate a new wave of genetic association studies aiming to identify rare, functional and causal variants for a broad range of complex traits and diseases.

As in conventional GWAS studies, associations identified in the UKB WES data will require stringent replication to minimize false discoveries, especially in rare variants orientated era. Unfortunately, for most studied traits, the community is still in short of adequate replication sample with sequenced genotype and matchable phenotype. On the other hand, the UKB has another ~300,000 participants who, despite not being released for their WES data yet, have as rich phenotypes as those WES participants have. This latter part of participants were assayed by genome-wide genotyping array (GWAS), and were imputed into the UK10K haplotype, 1000 Genomes project phase 3 and Haplotype Reference Consortium reference panels. Nonetheless, this imputed dataset is questionable for both coverage and imputation accuracy for rare coding variants [2].

In this study, we propose to utilize the combination of genetic data of UKB participants who were both sequenced and genotyped to impute sequenced variants of UKB participants who were genotyped only. We then propose to utilize the imputed participants as replication sample to replicate trait-associated LoF variants identified in the discovery WES sample. We will study genotype imputation accuracy for variants at a broad frequency spectrum, particularly for rare ones. As applications, we will conduct two-stage association studies of LoF variants for estimated heel bone mineral density (eBMD) and 4 lipid traits: high-density-lipoprotein cholesterol (HDL-C), low-density-lipoprotein cholesterol (LDL-C), triglycerides (TG) and total cholesterol (TC). We show that this WES based genotype imputation is accurate even for ultra-rare variants, therefore provides an efficient option for the identification of trait-associated LoF variants in the UKB cohort.

## Materials and Methods

### The UK Biobank cohort

The UK biobank (UKB) cohort is a large prospective cohort of ~500,000 participants from across the United Kingdom, aged between 48 and 73 at recruitment. Ethics approval for the UKB study was obtained from the North West Centre for Research Ethics Committee (11/NW/0382), and informed consent was provided by all participants. This study (application number 41542) was covered by general ethical approval for the UKB study.

### Genome-wide genotyping array data

Genome-wide genotyping (hereafter referred as GWAS) data is available for 488,000 UKB participants. In brief, genotype calling was performed by Affymetrix on two closely related purpose-designed arrays: ~50,000 participants were run on the UK BiLEVE Axiom array and the remaining ~450,000 were run on the UK Biobank Axiom array. The dataset combines results from both arrays and there are 784,256 autosome markers. In addition, the dataset was phased and imputed into UK10K haplotype, 1000 Genomes project phase 3 and Haplotype Reference Consortium reference panels. A total of ~92 million variants were generated by imputation.

### WES data

WES data is available for ~200,000 UKB participants [3]. In brief, the first tranche of WES was made available for 50,000 UK Biobank participants in March 2019, and the data for an additional 150,000 participants was made available in October 2020. The IDT xGen Exome Research Panel v1.0 probes target 39 Mbp of the human genome (19,396 genes). Multiplexed samples were sequenced with dual-indexed 75×75 bp paired-end reads on the Illumina NovaSeq 6000 platform.

The UKB 200K release was analyzed with an updated Functional Equivalence (FE) protocol [3] (referred to as the OQFE protocol). The OQFE protocol aligns and duplicate-marks all raw sequencing data (FASTQs) to the full GRCh38 reference in an alt-aware manner [4]. The OQFE CRAMs were then called for small variants with DeepVariant to generate per-sample gVCFs. These gVCFs were aggregated and joint-genotyped with GLnexus [5] to create a single multi-sample VCF (pVCF) for all UKB 200K samples. PLINK files were derived directly from this pVCF.

### Sample selection

The included participants were from the eligible genetically Caucasian (data field 22006) population, which was determined by a combination of self-reported “White British” and principal component derived genetic ancestry. As quality control (QC) procedure, participants who had a self-reported sex inconsistent with the genetic sex, whose sex chromosome was aneuploid, or who withdraw their consent were removed.

### Initial data QC

In the WES data, genetically duplicated participants were identified with the KING software with the “--duplicate" option [6]. Duplicated participants may represent either identical twins or repeat sampling of same participant, and were removed to avoid over-representation when forming reference panel for genotype imputation. At the variant level, variants having only one minor allele count (MAC) are prone to be produced due to sequencing error. In addition, they were not able to be evaluated for their imputation accuracy in subsequent genotype imputation. Therefore, they were removed from the WES data too. The following criteria were then applied to exclude WES participants and variants in PLINK [7]: individial-level genotype missingness>10%, variant-level genotype missingness>10%, or Hardy-weinberg equilibrium (HWE) P<1×10^−15^.

In the GWAS data, the following similar criteria were applied to exclude participants and variants: individual-level missingness>10%, variant-level genotype missingness >10%, or HWE P<1×10^−15^.

### Merging WES data and GWAS data

All WES participants have GWAS data too. The two datasets at these overlapping participants were merged into a combined reference data. Prior to merging, a series of filters were executed. We first converted the coordinates of the GWAS variants, which are based on the GRCH37 genome assembly, to the GRCH38 genome assembly coordinates. Variants failing to be converted were removed from the GWAS data. Excluded from the GWAS data also included palindromic variants (e.g., with A/T or G/C polymorphisms), whose strand could not be determined unambiguously. All alleles in both datasets were reported based on the forward genome orientation, which were double-checked by comparing the reported alleles with the forward-orientated reference genome sequence. Variants having inconsistent alleles were removed.

There are variants overlapping between the WES and the GWAS data. For these variants, we executed additional QC procedures. We first compared alleles and removed variants whose alleles are incompatible (e.g., A/G vs. A/C polymorphisms) from the GWAS data. We then calculated genotype discordance rate (GDR), which is defined as the proportion of genotypes that differ between the two datasets. Unusual GDR implies either sequencing error or genotying error. For sake of accuracy, they were removed from both datasets. The GDR exclusion threshold was set to be 5%.

After the above a series of QC filtering, variants from the WES data and the GWAS data were merged into a combined reference data containing overlapping participants. This dataset will be used as reference panel for subsequent genotype imputation. When merging overlapping variants, genotypes from the WES data were kept while those from the GWAS data were discarded.

### Reference data phasing

We next phased the reference data with SHAPEIT software [8]. To facilitate phasing computation, the whole genome was divided into dozens of genomic chunks each containing 10,000 target variants plus 500 kb buffer region to both directions whereas applicable. The length of chunks may vary depending on the local density of target variants.

### Reference data imputation

After the reference data was phased, it was then used as reference panel for imputation. We first imputed the reference data itself using a leave-100-out imputation strategy, for the purpose of evaluating imputation accuracy. Specifically, the participants in the reference data were randomly allocated into over one thousand non-overlapping sub-samples each containing 100 participants. In each sub-sample, genotypes from only the GWAS data (including overlapping variants) were retained, while the remaining variants from the WES data were masked and then imputed by the reference panel formed by haplotypes from all but the 100 to-be-imputed participants. The imputation was performed in each genomic chunk separately. Variants in buffer regions were ignored after imputation.

The imputation was done by FISH, a hidden Markov model (HMM)-based imputation method developed by us previously [9]. The following three features of FISH make its performance comparable to existing popular methods [10]: first, FISH calculates an exact form of HMM transition matrix in a linear computation complexity so the imputation accuracy is lossless; second, FISH models on a compact representation of reference haplotypes so the running time and memory usage could be largely improved. Third, FISH imputes directly diploid genotypes in a linear computation complexity so the test sample do not need to be phased *a prior*.

### Imputation accuracy measure

The imputed variants were evaluated for their imputation accuracy by comparing imputed genotypes with exome-sequenced genotypes in the reference data. Two accuracy measures were used: the first one is r^2^, which is defined as the correlation coefficient between true genotype and imputed hard-called genotype; the second one is GDR as defined above.

### Imputing the remaining GWAS sample

The remaining un-exome-sequenced GWAS participants were imputed by the above reference data with the same approach.

### Comparison with existing UKB imputed data

The UKB also released imputed genotypes by imputing the GWAS sample into the UK10K haplotype, 1000 Genomes project phase 3 and Haplotype Reference Consortium reference panels (hereafter called 1474 data). We compared the two imputed datasets in the overlapping participants. Comparison measures are again r^2^ and GDR.

### Loss-of-function variants annotation

Downstream association analyses were conducted at loss-of-function (LoF) variants only. Variants were annotated by VEP software [11]. Variants annotated as stop_gained, start_lost, splice_donor, splice_acceptor, stop_lost and frameshift were considered predicted LoF variants [2].

### Applications

As applications, we analyzed the association of LoF variants with eBMD and 4 plasma lipid traits in the UKB cohort: HDL-C, LDL-C, TG and TC.

For each trait, the total sample was divided into two sex groups. Within each group, phenotypic outliers were monitored and excluded by the Tukey fence inter-quantile range (IQR) approach [12]. Raw phenotypes from both groups were then combined together, followed by covariate adjustment by age, sex and the top 10 genetic PCs. The residuals after adjustment were normalized into inverse quantiles of standard normal distribution, which were used for subsequent association analysis.

### Association test in the discovery WES sample

The WES sample was used as the discovery sample. Two types of association were performed. The first one was the single variant based association test. The second one was gene-based burden test of rare LoF variants (MAF<1%). In the latter, all rare LoF variants within gene were collapsed into a single indicator of LoF carrier of that gene. Both single variant test and burden test were conducted by the linear mixed model with BOLT-LMM [13], in which the top 10 PCs were used as covariates.

### Replication in the WES-imputed GWAS data

The associated variants/genes were replicated in the imputed GWAS sample by single variant test or burden test. For burden test, only well imputed variants (r^2^>=0.8) were collapsed. Again, all associations were examined by the linear mixed model with BOLT-LMM in which the top 10 PCs were taken as covariates.

## Results

### Sample preparation

The WES data include 200,643 UKB participants, of whom 167,048 are eligible Caucasians. Forty-eight participants from 24 pairs were identified as duplicates and were excluded. Over 16 millions variants were called. After removing variants having only one MAC, 10,272,201 variants are retained. Further QC filtering on missingness and HWE retains 7,521,474 variants.

The GWAS data involve 408,915 Caucasian participants and 784,256 variants. QC filtering on missingness and HWE retains 408,911 participants and 689,184 variants. 531 variants failing to be converted to the GRCH38 assembly were filtered out. Another 41,911 palindromic variants were removed. In addition, 806 variants, though they were declared to be, are not on the forward orientation. These variants were removed from the GWAS data.

When comparing overlapping variants between the WES and the GWAS datasets, 1,840 variants are not allelic compatible, and were removed from the GWAS data. Additional 209 variants have unusually high GDR (>5%) between the two datasets, and were removed from both datasets.

The final clean data include 167,000 overlapping participants and 8,080,046 variants: 7,436,159 (92.0%) variants belong to the WES data only, 558,781 (6.9%) variants belong to the GWAS data only, and 85,106 (1.1%) variants overlap between the two datasets. Up to 91.6% (7,405,199) variants are rare ones (MAF<1%), while common (MAF>=5%) and less common (1%<=MAF<5%) variants account for only 5.1% (411,808) and 3.3% (263,039), respectively. These 167,000 overlapping participants and 8,080,046 variants were merged into one combined reference data.

The remaining 241,911 non-overlapping participants only have the GWAS data at 643,887 variants but not exome-sequenced. Their missing genotypes will be imputed by the merged reference data.

### Overall imputation accuracy

Using a leave-100-out imputation strategy, we divided the 167,000 participants of the reference data into 1,670 sub-samples each containing 100 participants. In each sub-sample, we masked genotypes at 7,436,159 exome-sequenced variants and then imputed them by the reference panel formed by the remaining 166,900 participants.

A total of 99,730 (1.34%) variants have no imputed polymorphism, all of them are rare (MAF<0.1%). For the remaining 7,336,429 variants, the average r^2^ and GDR are 0.58 and 0.04%, respectively. Below we will present r^2^ for illustration. In line with previous studies [14, 15], imputation accuracy increases with MAF (**Figure 1**). For 81,238 (1.11%) common variants, their missing genotypes were imputed with nearly perfect accuracy (r^2^=0.986). The average r^2^ for 46,405 (0.63%) less common variants gets slightly inferior, but still maintains at a high level (r^2^=0.952).

**Figure 1.**
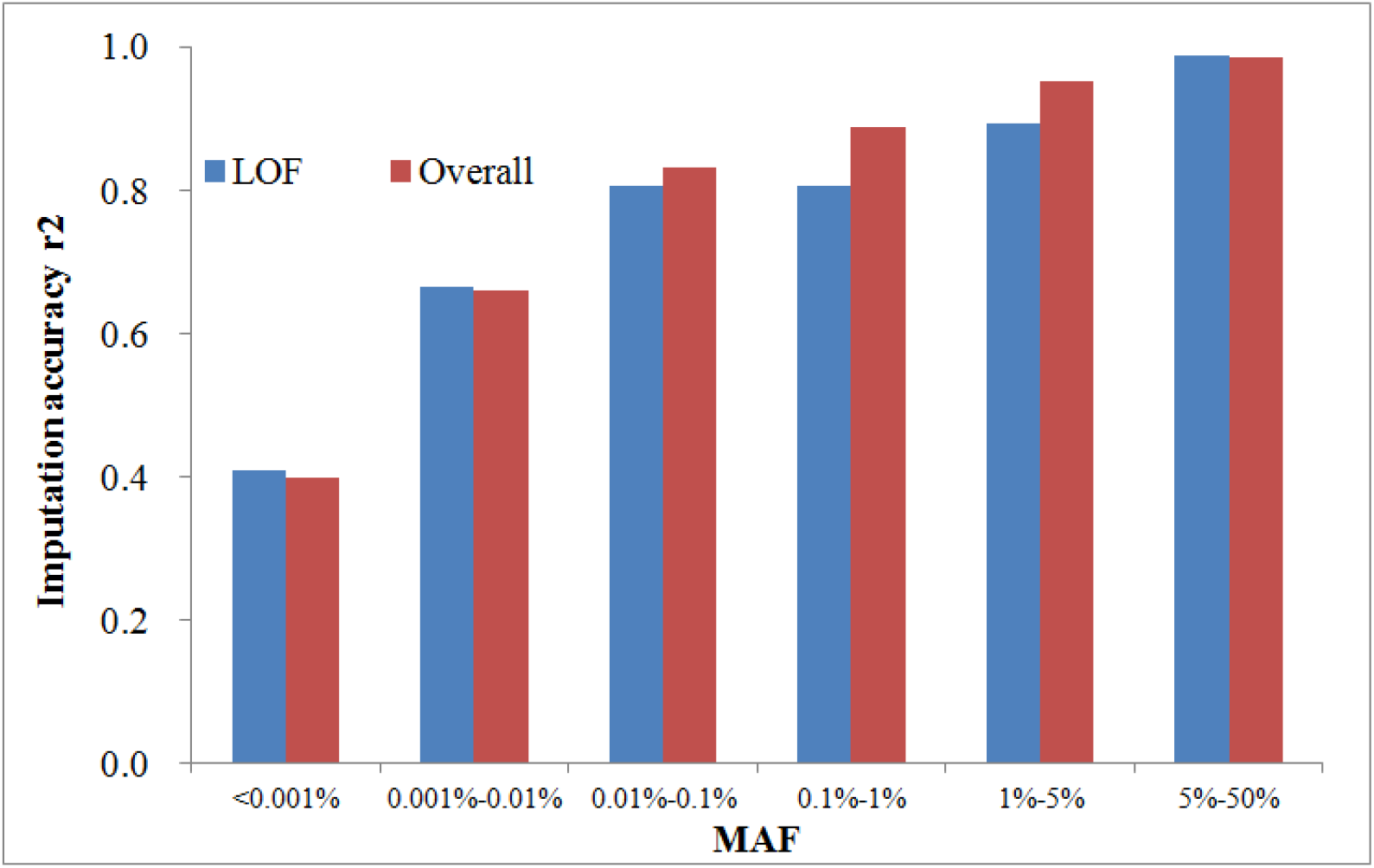
Imputation accuracy in the reference panel of 167,000 UKB Caucasian participants. The whole-exome sequencing (WES) genotypes and the genome-wide genotyping (GWAS) genotypes of 167,000 UKB Caucasian participants were merged into a combined reference panel, which was subjected haplotype phasing. Using a leave-100-out imputation strategy, the sample was randomly divided into 1,670 sub-samples each containing 100 participants. In each sub-sample, genotypes from only the GWAS data were retained, while the remaining variants from the WES data were masked and then imputed by the reference panel formed by haplotypes from all but the 100 to-be-imputed participants. Imputed genotypes were then compared with exome-sequenced variants to evaluate imputation accuracy, which was measured by squared correlation coefficient (r^2^) between the two sets of genotypes.

For a large number of rare variants, an inspiriting observation is that imputation accuracy still maintains at a modest to high level at all frequency ranges including ultra rare ones. For 117,202 (1.60%) variants falling in the MAF interval 0.1%-1.0%, the average r^2^ is as high as 0.887. When the interval gets down to 0.01%-0.1%, the 550,797 (7.51%) variants have an average r^2^ of 0.832. Next to it is the interval 0.001%-0.01%, the 3,571,280 variants in which take up the largest proportion (48.68%), and the average r^2^ is 0.659. Scaling to the total 167,000 participants, this interval corresponds to 4-34 MACs, a quite small number for such a large sample size. At last, at the smallest interval 0.001% or below, corresponding to only 2 or 3 MACs, the average r^2^ for 2,969,507 (40.48%) variants is still 0.400.

### LoF imputation accuracy

LoF variants are a class of variants truncating amino acid sequence, and are of primary interest. Of the above imputed variants, 139,848 are predicted to be LoF variants; nearly all of them (139,560, 99.8%) are rare ones. The average r^2^ and GDR for 288 (0.21%) common and less common LoF variants are 0.942 and 1.54%, respectively, which are slightly inferior to overall variants (0.974 and 1.39%). There are 593 (0.42%) LoF variants in the MAF interval 0.1%-1.0%, the average r^2^ of which is 0.807, again inferior to overall variants (0.887). Interestingly, when the MAF interval gets further down to 0.01%-0.1%, the average r^2^ of the 5,178 (3.70%) LoF variants changes little (0.805) to the variants in the 0.1%-1.0% interval, despite still being inferior to overall variants (0.832) in this interval. Variants in the 0.001%-0.01% interval account for 44.8% (62,689) of total LoF variants, with an average r^2^ of 0.664. At last, for the rarest 71,100 (50.8%) LoF variants whose MAF is below 0.001%, the average r^2^ is 0.410, nearly equivalent to that for overall variants.

By setting a threshold r^2^>=0.8 or r^2^<0.3 for well imputed or poorly imputed variants, we observed that 34.5% (48,261) of total LoF variants were well imputed. Apart from the 133,789 rarest variants with MAF below 0.01%, this proportion increases up to 73.7% (4,465/6,059). In contrast, 31.1% (43,530) LoF variants were poorly imputed; almost all of them (43,147, 99.1%) have a MAF<0.01%. The proportion of poorly imputed variants in the remaining 6,059 LoF variants is only 6.32%.

### Comparison with the UKB 500K imputed data

Of the above 139,848 imputed LoF variants, only 10,982 (7.85%) have same genomic coordinates as variants in the UKB released imputed data (hereafter referred as the “1474” data). Check of allele compatibility identified 6,808 compatible variants that are present in both imputed datasets. To avoid mis-strandness, palindromic variants were removed, leaving the final number of comparable variants being 6,080, including 113 common variants, 90 less common variants, 349 at MAF interval 0.1%-1%, 1,451 at 0.01%-0.1%, 2,913 at 0.001%-0.01%, and 1164 at 0.001% or below.

For WES imputed data, the imputation accuracy at these overlapping variants is similar to that in overall variants at all MAF intervals. On the other hand, the 1474 data have largely reduced imputation accuracy at all MAF intervals (**Figure 2**). Of the 6,080 compared variants, the number of well imputed variants in the WES-imputed data is 4,935, while that in the 1474 data is only 549, or a nine-fold reduction. Taking the total LoF variants into account, the present WES-imputed data not only increases the number of well imputed LoF variants by 87-fold (48261/549), but also have largely increased imputation accuracy in all MAF intervals including common and rare variants.

**Figure 2.**
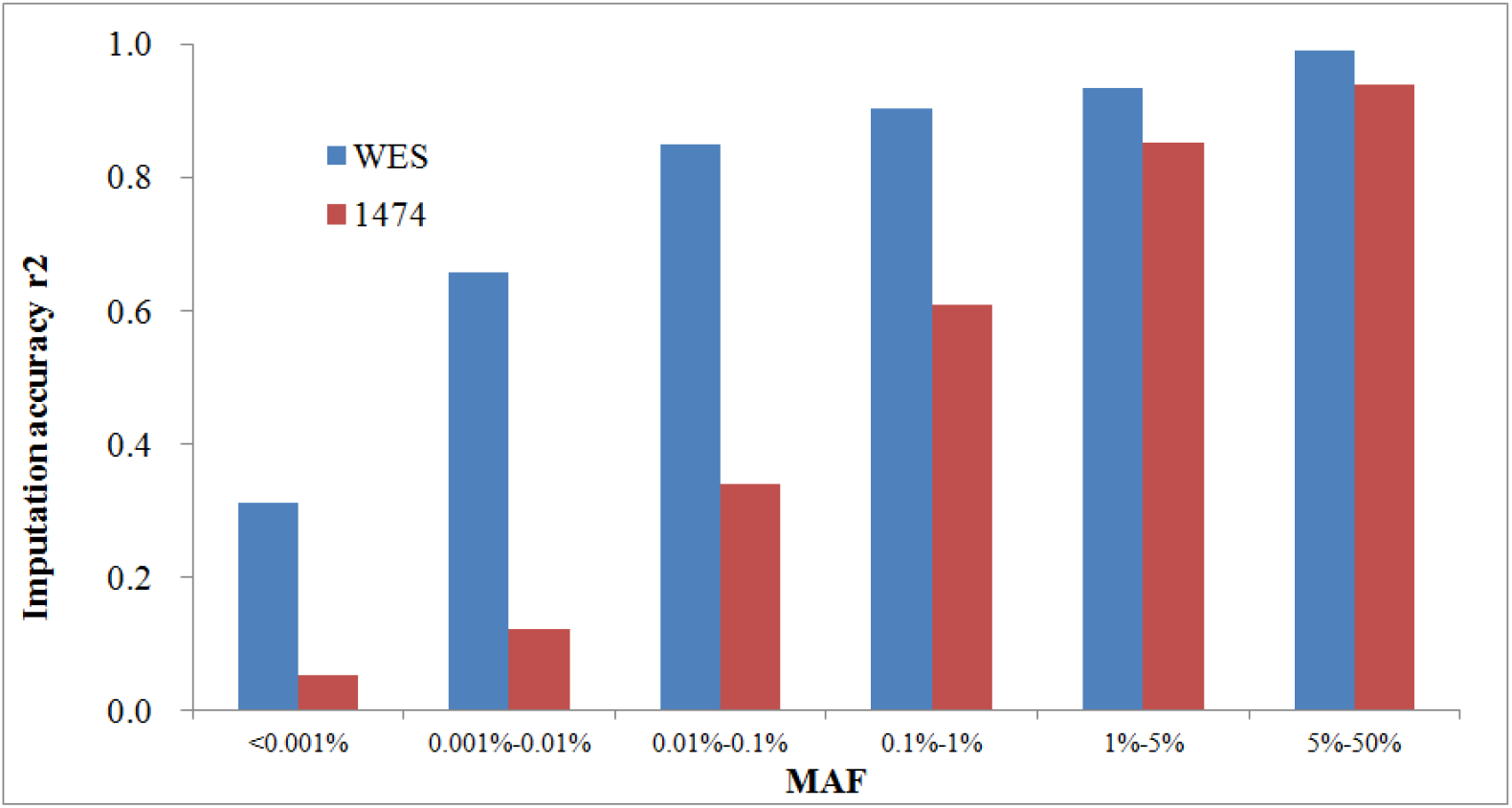
LoF imputation accuracy comparison with existing imputed genotypes. A total of 6,080 LoF variants are present in both the present WES-imputed genotypes (WES) and the UKB released imputed genotypes (“1474” dataset). Their imputation accuracy (r^2^) was evaluated and compared on the 167,000 sequenced participants whose genotypes were known.

### eBMD association study

We first analyzed eBMD association. At the Bonferroni-corrected significance level P<3.13×10^−7^ (0.05/159,818), single variant test of the discovery sample identified one rare LoF variant 4:87845484:D:1 (rs778732516, MAF=0.13%, BETA=−0.43, P=2.20×10^−12^) in *MEPE* gene. Gene-based burden test identified the same gene *MEPE* (BETA=−0.37, SE=0.04, P=4.00×10^−12^) at the gene significance level P<3.06×10^−6^ (0.05/16,325), where the burden statistic collapses 19 rare LoF variants with a cumulative MAF of 0.21%. No other single variant or gene is significant at the specified significance levels.

We next replicated the top signals in the WES-imputed GWAS sample. For gene-based test, we sought to replicate the top 10 genes, whose discovery p-values range from 7.24×10^−4^ to 2.20×10^−12^. Single variant test successfully replicates 4:87845484:D:1 with very strong signal (BETA=−0.35, P=1.40×10^−12^). Gene-based burden test replicates *MEPE* as well (BETA=−0.34, P=9.30×10^−19^), but none of the other 9 genes could be replicated at the desired significance level P<0.005 (0.05/10).

Of note, though the discovery burden signal of *MEPE* is mainly driven by 4:87845484:D:1, sensitivity analysis shows that the burden signal excluding 4:87845484:D:1 remains nominally significant (BETA=−0.23, P=3.40×10^−3^), implying the presence of additional responsible variant. Scrutiny of individual variant associations highlights another variant 4:87845066:D:4, which is nominally significant and is rarer than 4:87845484:D:1 (MAF=0.05%, BETA=−0.40, P=1.70×10^−4^). Further sensitivity analysis excluding 4:87845484:D:1 and 4:87845066:D:4 becomes non-significant (P=0.68), implying no additional association source. This observation is strengthened in the replication sample by both extremely significant signals at both variants (4:87845484:D:1, BETA=−0.35, P=1.40×10^−12^; 4:87845066:D:4, BETA=−0.53, P=6.30×10^−13^) and the fact that neither any other single variant nor the burden test excluding both variants is nominally significant.

Multiple variants around *MEPE* were previously reported to be associated with BMD and fracture risk [16-18]. In a recent WES study of the same eBMD trait in the smaller ~50,000 UKB participants [2], both 4:87845484:D:1 (P=6.56×10^−4^) and 4:87845066:D:4 (P=2.19×10^−3^) were nominally significant. A third variant 4:87844983:D:1 was nominally significant too (P=2.79×10^−3^) in that study. In the present study, however, 4:87844983:D:1 is significant in neither the discovery (P=0.17) nor the replication sample (P=0.63).

Both 4:87845484:D:1 (MAF=0.13%) and 4:87845066:D:4 (MAF=0.05%) are frameshift deletion resulting in truncated protein at the 237^th^ and 292^th^ amino acid, respectively. Both variants were predicted by VEP to have high impact, implying their disrupting effect to the *MEPE* structure and function. The protein encoded by *MEPE* plays an important role in osteocyte differentiation and bone homeostasis [19].

### Lipids association study

As a second application, we analyzed 4 lipid traits, namely HDL-C, LDL-C, TG and TC. At the same single variant significance level P<3.13×10^−7^, the discovery sample identified 65 associations from 36 LoF variants from 22 genes for one or more lipid traits. Of them, 13 are common variants while the other 23 are rare ones, with MAF ranging from as low as 6.6×10^−4^% to 49.7%. We then sought to replicate these variants in the imputed GWAS sample. Up to 30 of the above 36 variants were genotyped or well-imputed (r^2^>=0.8). In the replication sample, 56 (86.2%) of the 65 associations are successfully replicated at stringent significance level P<7.69×10^−4^ (0.05/65), all with consistent effect direction. Of the remaining 9 non-replicated associations from 7 variants, 2 variants had no imputed polymorphism due to extremely low frequency (MAF<1.0×10^−3^%); 2 were poorly imputed (r^2^=0.00 and 0.13); 3 are nominally significant but the signals do not achieve the specified significance level (P=8.60×10^−3^-1.50×10^−3^, **Supplemental table 1**). Thirty-three of the above 56 identified associations were driven by rare variants (MAF<1%). Interestingly, in all cases, rare alleles identified for HDL-C tend to increase HDL-C level while those identified for the other three traits tend to decrease their levels.

Gene-based burden test of rare LoF variants identified 22 significant associations in the discovery WES sample at the gene significance level P<3.06×10^−6^. As in the case of eBMD, we further expanded the genes to be replicated to the top 10 hits of each trait, and set the replication significance threshold to 1.25×10^−3^ (0.05/10/4). Seven-teen of the above 22 associations are successfully replicated. Three additional associations achieve P<1.25×10^−3^ in the replication sample, increasing the total number of successfully replicated gene associations to be 20 (17+3, **Supplemental table 2**).

Together, individual variant test identified and replicated 56 associations from 39 gene-trait pairs, and gene burden test identified and replicated 20 pairs of association. Twelve pairs overlap between both tests, leaving the total number of identified association pairs to be 47. They are from 26 genes in total. We defined a pair of gene-trait association to be novel if the gene is located at least 1 Mb from signals reported by two previous large-scale studies [20, 21] for that particular trait. With this definition, 3 gene-trait pairs are found to be novel, 2 genes *PLIN1* and *ANGPTL3* for HDL and one gene *PDE3B* for TG (**Table 1**).

**Table 1.**
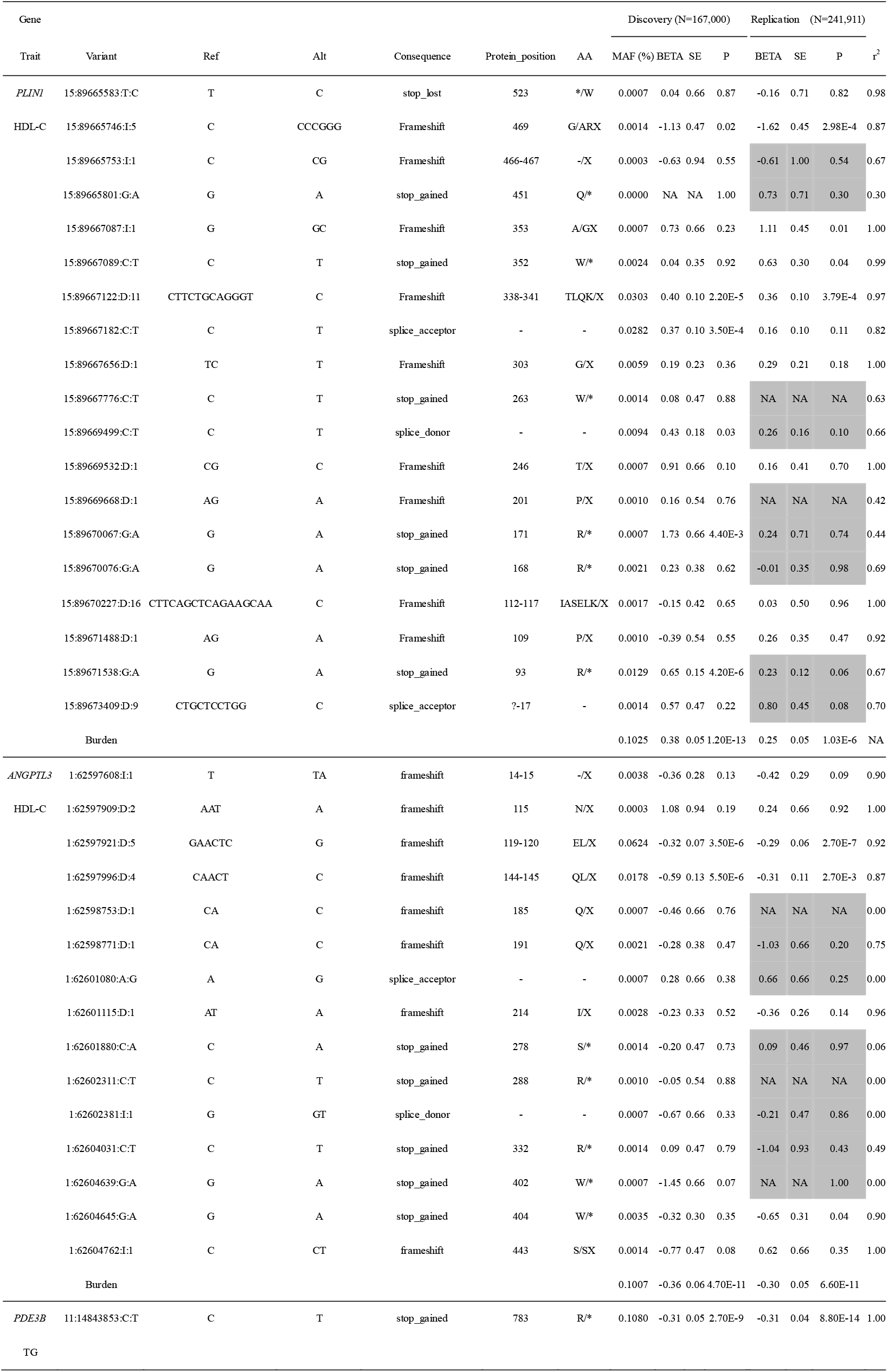
Novel gene-trait associations identified in the present study. Notes: Ref, reference allele; Alt, alternative allele; Protein_position, the protein position in which the mutation occurs; MAF, minor allele frequency, or cumulative minor allele frequency in case of burden test; BETA, regression coefficient; SE, standard error of BETA; P, p-value; r^2^, imputation correlation coefficient. Variants marked in gray were not collapsed due to r^2^<0.8.

Both *PLIN1* and *ANGPTL3* were identified by gene burden test. *PLIN1* contains 19 rare LoF variants, 10 of which were well-imputed (r^2^>=0.8). No single variant achieves the preset significance level in the discovery sample. In contrast, the aggregate signal of multiple variants gets extremely significant (BETA=0.38, P=1.20×10^−13^). The replication sample collapses the 10 well-imputed LoF variants and replicates the association (BETA=0.27, P=6.30×10^−9^). *PLIN1* encodes a protein that coats lipid storage droplets in adipocytes, thereby protects them until they are broken down by hormone-sensitive lipase [22]. *ANGPTL3* contains 15 rare LoF variants, 7 of which were well imputed. The aggregate signal in the discovery sample is mainly driven by 2 variants 1:62597921:D:5 (P=3.5×10^−6^) and 1:62597996:D:4 (P=5.5×10^−6^), both of which are significant in the replication sample too (P=2.7×10^−7^ and 2.7×10^−3^). 1:62597921:D:5 is also significant with all other 3 lipid traits in single variant test, implying the pleiotropic effect of this gene to all 4 studied lipid traits. At last, the association of *PDE3B* with TRI was identified by a single LoF variant 11:14843853:C:T (MAF=0.11%, discovery BETA=−0.31, P=2.70×10^−9^; replication BETA=−0.31, P=8.80×10^−14^).

### Allelic heterogeneity

Of the identified 20 gene-trait burden associations, 10 contain variants that are already individually significant at P<3.13×10^−7^ in the discovery sample. Sensitivity analysis after excluding these variants from collapsing remains nominally significant (P<0.05) at all associations (**Supplemental table 3**). For the other 10 gene associations that contain no extremely significant variants, leave-one-out sensitivity analysis remains nominally significant after excluding any one single variant. Altogether, the above results suggest that the aggregate signal is driven by multiple variants for any identified gene association, demonstrating the presence of allelic heterogeneity [23].

### Linkage disequilibrium with known GWAS loci

Twenty-three of the above 56 single variant associations are from 13 common variants (MAF>=5%). Of note, all but one of them are in strong linkage disequilibrium (LD, r^2^=0.75-1.00) with one or more common variants that were previously reported, where the LD pattern for the remaining one variant 6:160139865:D:8 is weaker (LD r^2^=0.20 with rs1888727). Conditioning on each of the 13 LoF variants, the signals at corresponding neighbor variants get weaker or even vanished.

In contrast, for the 33 associations carried by 17 rare LoF variants (MAF<1%), except for 2 variants which themselves were reported by previous studies, none of the other variants are in LD with previous reported variants (r^2^<0.05), implying these variants representing additional association sources.

## Discussion

In this study, we have proposed to combine the WES data and the GWAS data of the ~200,000 UKB participants to impute the remaining UKB participants who had the GWAS data only, and to use the imputed data as the replication of association identified in the discovery WES sample. We have empirically studied imputation accuracy at full frequency spectrum and applied to association studies of LoF variants with eBMD and 4 lipid traits. The results showed that the imputation accuracy is modest to high at the vast majority of variants including ultra-rare ones. The imputed sample well-replicated the findings discovered in the discovery WES sample, providing solid confidence towards true associations.

During the time of the preparation of this manuscript, Barton et al. published a similar UKB exome-sequencing imputation study [24]. The major difference between the two studies is the use of first tranche of ~50,000 WES participants as reference panel in that study versus the use of all ~200,000 WES participants in the present study. This difference could make the present study advantageous in two aspects: first, it imputed more coding variants, as the number of sequenced variants from the ~200,000 participants is larger than those from the ~50,000 participants [3]. Second, using a reference sample with enlarged size by up to 4-fold, imputation accuracy is expected to be higher in the present study.

An impressive observation from this study was the ability to identify replicable associated variants at extremely low frequency. One example is the association of 2:21009310:G:A (discovery BETA=−2.43, P=1.60×10^−7^) with LDL-C. This variant’s MAF is only 1.27×10^−3^%, corresponding to 4 MACs in the discovery 167,000 participants. The fact that this variant is successfully replicated (replication BETA=−2.18, P=9.00×10^−5^) strengthens the evidence towards its true association. Certainly, this ability relies on variant effect size. In this example, the estimated large per allele effect of −2.18 by the replication sample corresponds to 26.6% statistical power for a discovery sample of 167,000 participants at the specified significance level 3.13×10^−7^. The replication sample of over 200,000 participants has a nearly perfect statistical power of 93.9% to replicate it at the significance level 7.69×10^−4^. Despite being imperfectly imputed (r^2^=0.39), the power gained in the replication sample is still considerable. Identification of such LoF variants has important clinical application, as they are potential drug target to treat disease.

Another observation worth of mention is that even in a likely causal gene, not all LoF variants are associated with the target trait. The association of *MEPE* gene with eBMD, for example, is driven by only 2 variants although the aggregate signal collapsed 19 LoF variants. Their significance was verified in not only the discovery sample but also the replication sample. Although the non-significance at the other variants could be attributable to insufficient statistical power or LoF variant annotation artifact [1], their null effect is the most likely reason. The mechanism behind this phenomenon deserves further investigation.

The exomes of all ~500,000 UKB participants are scheduled to be released in the near future. Before then, the data imputed in the present study will be valuable in exploring LoF variants associated with diseases as well as refining in previously reported loci. Another implication is the value of the use of WES sample to impute rare coding variants in existing GWAS data. With the emerging of open access ultra-large WES datasets such as the UKB WES sample, there will be an opportunity to impute numerous previous GWAS samples and study the role of rare coding variants in disease pathogenesis in these GWAS samples.

## Acknowledgements

This research was conducted using the UK Biobank resource under application number 41542. LZ and YFP were partially supported by the funding from national natural science foundation of China (31771417 to YFP) and by a project funded by the Priority Academic Program Development (PAPD) of Jiangsu higher education institutions (to LZ). The numerical calculations in this paper have been done on the supercomputing system of the National Supercomputing Center in Changsha.

## References

1. Karczewski, K.J., et al., The mutational constraint spectrum quantified from variation in 141,456 humans. Nature, 2020. 581(7809): p. 434–+.

2. Van Hout, C.V., et al., Exome sequencing and characterization of 49,960 individuals in the UK Biobank. Nature, 2020. 586(7831): p. 749–+.

3. Szustakowski, J.D., et al., Advancing human genetics research and drug discovery through exome sequencing of the UK Biobank. Nature Genetics, 2021. 53(7): p. 942–948.

4. Regier, A.A., et al., Functional equivalence of genome sequencing analysis pipelines enables harmonized variant calling across human genetics projects. Nature Communications, 2018. 9.

5. Van Hout, C., et al., Whole exome sequencing and characterization of coding variation in 49,960 individuals in the UK Biobank. European Journal of Human Genetics, 2019. 27: p. 1166–1167.

6. Manichaikul, A., et al., Robust relationship inference in genome-wide association studies. Bioinformatics, 2010. 26(22): p. 2867–73.

7. Purcell, S., et al., PLINK: a tool set for whole-genome association and population-based linkage analyses. Am J Hum Genet, 2007. 81(3): p. 559–75.

8. Delaneau, O., J. Marchini, and J.F. Zagury, A linear complexity phasing method for thousands of genomes. Nature Methods, 2012. 9(2): p. 179–181.

9. Zhang, L., et al., FISH: fast and accurate diploid genotype imputation via segmental hidden Markov model. Bioinformatics, 2014. 30(13): p. 1876–1883.

10. Howie, B., et al., Fast and accurate genotype imputation in genome-wide association studies through pre-phasing. Nat Genet, 2012. 44(8): p. 955–+.

11. McLaren, W., et al., The Ensembl Variant Effect Predictor. Genome Biol, 2016. 17.

12. Hoaglin, D.C., John W. Tukey and data analysis. Statistical Science, 2003. 18(3): p. 311–318.

13. Loh, P.R., et al., Efficient Bayesian mixed-model analysis increases association power in large cohorts. Nature Genetics, 2015. 47(3): p. 284–+.

14. Howie, B., et al., Fast and accurate genotype imputation in genome-wide association studies through pre-phasing. Nature Genetics, 2012. 44(8): p. 955–+.

15. Das, S., G.R. Abecasis, and B.L. Browning, Genotype Imputation from Large Reference Panels. Annual Review of Genomics and Human Genetics, Vol 19, 2018. 19: p. 73–96.

16. Kemp, J.P., et al., Identification of 153 new loci associated with heel bone mineral density and functional involvement of GPC6 in osteoporosis. Nature Genetics, 2017. 49(10): p. 1468–+.

17. Medina-Gomez, C., et al., Life-Course Genome-wide Association Study Meta-analysis of Total Body BMD and Assessment of Age-Specific Effects. American Journal of Human Genetics, 2018. 102(1): p. 88–102.

18. Zheng, H.F., et al., Whole-genome sequencing identifies EN1 as a determinant of bone density and fracture. Nature, 2015. 526(7571): p. 112–+.

19. Zelenchuk, L.V., A.M. Hedge, and P.S.N. Rowe, Age dependent regulation of bone-mass and renal function by the MEPE ASARM-motif. Bone, 2015. 79: p. 131–142.

20. Liu, D.J., et al., Exome-wide association study of plasma lipids in > 300,000 individuals. Nature Genetics, 2017. 49(12): p. 1758–+.

21. Willer, C.J., et al., Discovery and refinement of loci associated with lipid levels. Nature Genetics, 2013. 45(11): p. 1274–+.

22. Kimmel, A.R., et al., Adoption of PERILIPIN as a unifying nomenclature for the mammalian PAT-family of intracellular lipid storage droplet proteins. Journal of Lipid Research, 2010. 51(3): p. 468–471.

23. Hormozdiari, F., et al., Widespread Allelic Heterogeneity in Complex Traits. American Journal of Human Genetics, 2017. 100(5): p. 789–802.

24. Barton, A.R., et al., Whole-exome imputation within UK Biobank powers rare coding variant association and fine-mapping analyses. Nature Genetics, 2021.

